# Reply to: The pitfalls of negative data bias for the T-cell epitope specificity challenge

**DOI:** 10.1101/2023.04.07.535967

**Authors:** Yicheng Gao, Yuli Gao, Kejing Dong, Siqi Wu, Qi Liu

## Abstract

Predicting and identifying TCR-antigen pairings accurately presents a significant computational challenge within the field of immunology. The negative sampling issue is important T-cell specificity modeling and it is known clearly by the community that different negative data sampling strategy will influence the prediction results. Therefore, proper negative data sampling strategy should be carefully selected, **<u>and this is exactly what PanPep has noticed, emphasized and performed</u>**. Now we would like to clarify this point further by formulating this problem as a PU learning. Our findings suggest that the reshuffling strategy may generate potential false negative samples, which can adversely affect model training and result in biased model testing for PanPep. Furthermore, a proper comparison between different negative sampling strategies should be performed **<u>in a consistent way</u>** to make a proper conclusion. Finally, future updating to explore more possible and suitable negative sampling strategy is expected.

## Main article

We noticed that recently Pieter Meysman et al indicated the negative data sampling issue in T-cell epitope specificity prediction^1^. In light of the limited data available in this area, the negative sampling issue is generally important for biological data modeling, since biological experimental tests intend to record the positive results while ignore the negative results. We appreciated their efforts to raise this point for T-cell epitope specificity modeling, which is known clearly by the community that different negative data sampling strategy will influence the prediction results^2, 3^. Therefore, proper negative data sampling strategy should be carefully selected, **<u>and this is exactly what PanPep has noticed, emphasized and performed</u>**^4^. In short, as for the two commonly used negative sampling strategy, i.e., reshuffling based on positive pairs (**first strategy**) and randomly drawing from background repertories (**second strategy)**, **<u>PanPep prefers to select the second strategy</u>**, and the rational has been clearly indicated in the manuscript. Now we would like to clarify this point further by formulating this problem as a PU learning and calling for more attentions on this point.

1. In the current stage of the study, actually the computational essence of TCR-epitope recognition problem can be formulated as Positive & Unlabeled Data Learning (PU Learning) in terms of the lack of clearly defined negative samples^5^. PU Learning involves having only positive data and some unlabeled data, for which the negative status is unknown (Fig 1). Therefore, generating proper negative data is important. **<u>This is also well known for other communities besides</u> TCR-peptide recognition, such as for protein-ligand binding prediction**^**6**^ **etc**. However, in the TCR-peptide recognition community, it is well known that the cross-reactivity of TCRs is a significant challenge in the TCR-peptide recognition problem and it can pose difficulties for many models in the pre-processing stage^2^. Experimental methods can have a high false-negative rate^1^, resulting in many potential cross-reactivity in existing known binding TCRs, making PU Learning more challenging. In the absence of experimental negatives, two main negative sampling strategies have emerged, including reshuffling based on positive pairs (**first strategy**) and randomly drawing from background repertories (**second strategy**). The first strategy assumes that TCRs have no cross-reactivity, while the second strategy assumes that TCRs from background repertories do not react to the epitopes of interest^2^. Previous mainstream models have been built based on either of these two assumptions^7-10^. **<u>While it should be noted that the training and testing should be kept with the same assumption to make an objective conclusion. The two tests performed by Pieter Meysman et al for PanPep is improperly built on an inconsistent assumption, i.e., training it on the second strategy while testing it on the first strategy, therefore resulting in improper conclusions. In this case, the performance decrease of PanPep in the reshuffling test is SIMPLY due to the inconsistent assumption during the training and testing. So if you want to evaluate the model performance properly under two different negative sampling strategies, please do keep it in a consistent scenario for model training and testing</u>**.

**Fig 1.**
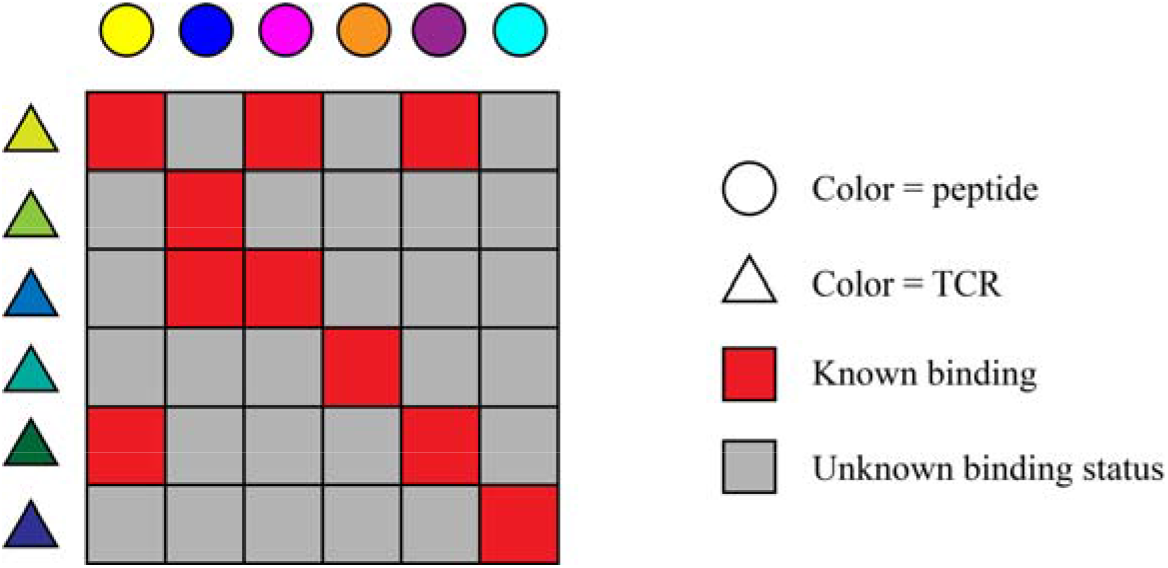
PU Learning problem in TCR-peptide recognition

**<u>2. The careful selection of proper negative sampling strategy is exactly what PanPep has considered. PanPep was built based on the second strategy due to its use of few-shot learning and the need for high-quality samples</u>**. This point has been clearly explained in the manuscript, i.e. “Since one TCR sequence might bind to different peptides with cross-reactivity and the meta learning module of PanPep is highly sensitive to the data quality, we must reduce the bias from the mislabels among the nonbinding TCRs as much as possible.” and “Considering the large number of TCRs in this healthy repertoire, randomly sampling a part of TCRs from this large pool as a control set has **<u>a very low probability</u>** of encountering TCRs binding to the given peptide.”

3. In order to present a more concretely and comprehensively analysis to support our selection, now we analyzed the cross-reactivity of TCRs in both the meta-dataset and the zero-dataset. In the meta-dataset, over 1,600 known TCRs among 29,057 unique TCRs exhibited cross-reactivity, and around 21.4% of TCRs (100+) among 543 unique TCRs in the zero-dataset had cross-reactivity (Fig 2), indicating the substantially existed cross-reactivity. We argued that using the reshuffling strategy in these datasets would bias the dataset and increase the likelihood of mislabeling negative samples compared to the second strategy.

**Fig 2.**
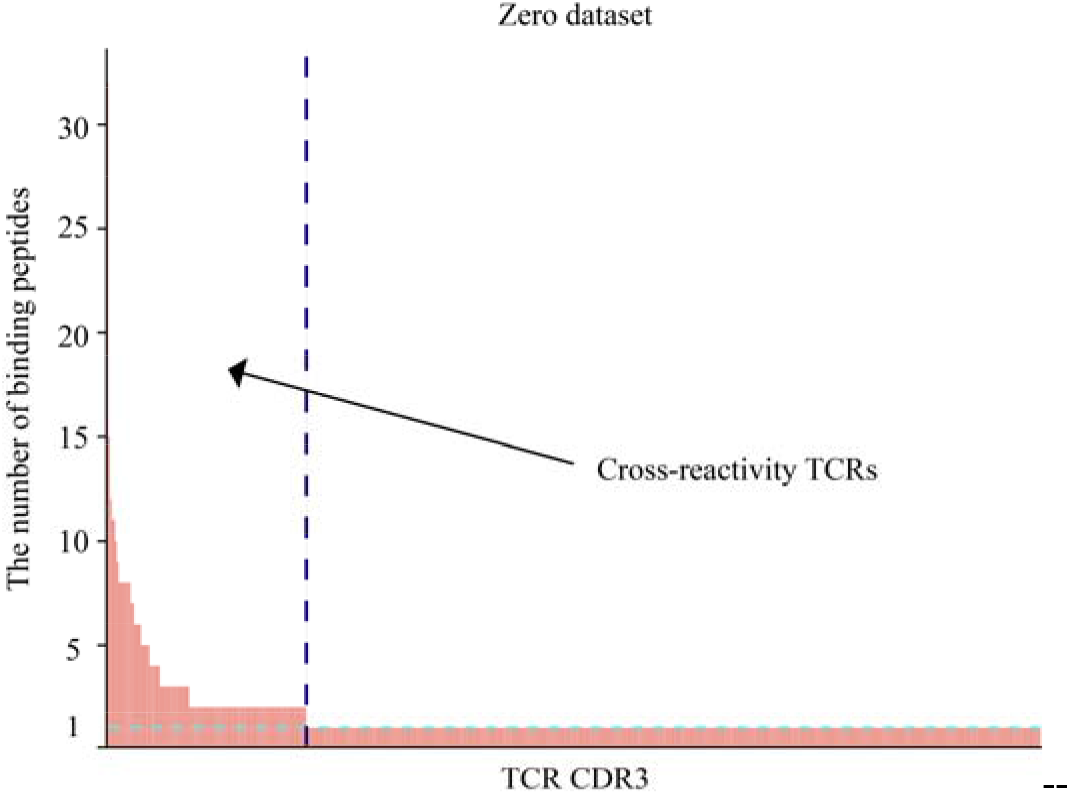
The case of cross-reactivity of TCRs in the zero-dataset.

To this end, we conducted the experiments to test the performance of PanPep where PanPep was trained on both two strategies and tested on both strategies (**2*2 training-testing**). Firstly, our original PanPep model, trained on the second strategy, achieved a ROC-AUC and PR-AUC of 0.708 and 0.715 in zero-shot testing using the second strategy, respectively (Table 1). **<u>This is what we exactly applied in PanPep. It can be seen that among the 4 scenarios, PanPep achieved the best in this case, indicating that the second strategy is proper for PanPep</u>**. In the testing with first strategy, where we reshuffled the zero-shot dataset 10 times, the model showed an average ROC-AUC and PR-AUC of 0.553 and 0.563, respectively (Table 1). Additionally, we trained a new PanPep model based on the first strategy and tested it on both strategies. This model achieved a ROC-AUC and PR-AUC of 0.55 and 0.56 in zero-shot testing with the first strategy, and 0.627 and 0.640 in zero-shot testing with the second strategy (Table 1). Notably, the background TCRs were not used in the training process. However, the new PanPep also showed the reduced performance when testing on the zero-shot dataset with the first strategy, potentially due to the cross-reactivity in the testing dataset, leading to mislabeled negative samples. In sum, our comprehensive tests demonstrating that **<u>(1) When testing PanPep in the secondary strategy, no matter what kinds of negative sample strategy has been selected in the training, PanPep both have certain prediction ability in the zero-shot testing (∼0.7 for training with the secondary strategy and ∼0.6 for training with the first strategy), indicating PanPep is able to solve the challenging zero-shot learning problem with certain prediction ability for both negative sampling strategies applied in the training; (2) Compared to the secondary strategy, training PanPep based on the reshuffling strategy would negatively impact the performance due to relatively low negative sample quality; Collectively, in the current study, we do not recommend to use reshuffling strategy, either for model training or computational testing</u>**.

**Table 1.** The performance of PanPep on zero-shot testing with different negative sampling strategies.

| Testing (Zero-shot) | Training (first strategy) | Training (second strategy) |
| --- | --- | --- |
| First strategy (ROC-AUC) | 0.551±0.004 | 0.553±0.005 |
| Second strategy (ROC-AUC) | 0.627 | 0.708 |
| First strategy (PR-AUC) | 0.556±0.005 | 0.563±0.005 |
| Second strategy (PR-AUC) | 0.640 | 0.715 |

4. Although the current version of PanPep applied the second strategy, future updating to explore more possible and suitable negative sampling strategy is expected. It should be noted that distinguishing binding TCRs from the background in the zero-shot scenario is very challenging. Our comprehensive tests have clearly indicated that existing tools failed to distinguish positive TCRs from background TCRs for zero-shot learning. The zero-shot learning for T-cell specificity modeling is challenging, seen as the ‘holy grail’ of immunology as indicated^11^. Now we would like to emphasize again the novelty of PanPep in addressing such zero-shot prediction facing the very challenging “long-tail” issue^12^, and this conceptual methodology novelty should not be overlooked.

In conclusion, (1) we appreciated the efforts and the comments raised here, while we would like to emphasize again the novelty of PanPep in addressing zero-shot prediction for T-cell epitope specificity prediction; (2) Also we would like to emphasize that the proper selection of negative sampling strategy has been carefully considered in PanPep. And PanPep prefers to use the second strategy considering the requirement of high quality data to train a meta learning model; (3) A proper comparison between different negative sampling strategies should be performed **<u>in a consistent way</u>** for training and testing; (4) PanPep can be directly applied for peptide-TCR binding prediction in a realistic scenario, where negative sampling is not required, while the experimental validation are expected. The negative sampling is only required for model training and computational evaluation, nevertheless, proper evaluating and comparing the model performance under different negative sampling strategies do require to keep the training and testing in a consistent way; (5) We expect that more data and an unbiased benchmarking experimental data could be generated and developed. More available data, including a more diverse set of available peptides, will certainly enhance the performance and generalization of PanPep; (6) Finally, more possible evaluation measurements and negative sampling strategies for T-cell epitope specificity prediction are expected in such negative sample absent scenario.

## Acknowledgement

This work was supported by the National Key Research and Development Program of China (Grant No. 2021YFF1201200, No. 2021YFF1200900), National Natural Science Foundation of China (Grant No. 31970638, 61572361), Shanghai Natural Science Foundation Program (Grant No. 17ZR1449400), Shanghai Artificial Intelligence Technology Standard Project (Grant No. 19DZ2200900), Shanghai Shuguang scholars project, Program of Shanghai Academic/Technology Research Leader, WeBank scholars project and Fundamental Research Funds for the Central Universities.

## Reference

1. Dens, C., Laukens, K., Bittremieux, W. & Meysman, P. The pitfalls of negative data bias for the T-cell epitope specificity challenge. Preprint at bioRxiv https://doi.org/10.1101/2023.04.06.535863 (2023).

2. Hudson, D., Fernandes, R.A., Basham, M., Ogg, G. & Koohy, H. Can we predict T cell specificity with digital biology and machine learning? Nature Reviews Immunology, 1–11 (2023).

3. Jiang, Y., Huo, M. & Cheng Li, S. TEINet: a deep learning framework for prediction of TCR–epitope binding specificity. Briefings in Bioinformatics 24, bbad086 (2023).

4. Gao, Y. et al. Pan-Peptide Meta Learning for T-cell receptor–antigen binding recognition. Nature Machine Intelligence, 1–14 (2023).

5. Elkan, C. & Noto, K. in Proceedings of the 14th ACM SIGKDD international conference on Knowledge discovery and data mining 213–220 (2008).

6. Chen, L. et al. Hidden bias in the DUD-E dataset leads to misleading performance of deep learning in structure-based virtual screening. PloS one 14, e0220113 (2019).

7. Lu, T. et al. Deep learning-based prediction of the T cell receptor–antigen binding specificity. Nature Machine Intelligence 3, 864–875 (2021).

8. Springer, I., Tickotsky, N. & Louzoun, Y. Contribution of t cell receptor alpha and beta cdr3, mhc typing, v and j genes to peptide binding prediction. Frontiers in immunology 12 (2021).

9. Luu, A.M., Leistico, J.R., Miller, T., Kim, S. & Song, J.S. Predicting TCR-epitope binding specificity using deep metric learning and multimodal learning. Genes 12, 572 (2021).

10. Gielis, S. et al. Detection of enriched T cell epitope specificity in full T cell receptor sequence repertoires. Frontiers in immunology, 2820 (2019).

11. Moris, P. et al. Current challenges for unseen-epitope TCR interaction prediction and a new perspective derived from image classification. Briefings in Bioinformatics 22, bbaa318 (2021).

12. Wang, D., He, F., Yu, Y. & Xu, D. Meta-learning for T cell receptor binding specificity and beyond. Nature Machine Intelligence, 1–3 (2023).

